# Time–space signatures of hybrid search resolution using EEG and eye movements concurrent recordings

**DOI:** 10.64898/2026.06.22.733836

**Authors:** Damián Care, Joaquín E. González, Matias J. Ison, Juan E. Kamienkowski

**Affiliations:** Laboratorio de Inteligencia Artificial Aplicada, Instituto de Ciencias de la Computación (Consejo Nacional de Investigaciones Científicas y Técnicas - Facultad de Ciencias Exactas y Naturales, Universidad de Buenos Aires), Argentina; Departamento de Física, Facultad de Ciencias Exactas y Naturales, Universidad de Buenos Aires, Argentina; School of Psychology, University of Nottingham, United Kingdom; Departamento de Computación, Facultad de Ciencias Exactas y Naturales, Universidad de Buenos Aires, Argentina; Maestría de Explotación de Datos y Descubrimiento del Conocimiento, Facultad de Ciencias Exactas y Naturales, Universidad de Buenos Aires, Argentina

**Keywords:** Hybrid Search, EEG and eye movements concurrent recordings, deconvolution models

## Abstract

Understanding how the brain supports visual search in naturalistic environments—where attention and memory must work together to find targets among distractors—requires analysing neural signals where responses overlap in time and multiple environmental variables simultaneously interact. Conventional event-related methods cannot disentangle these overlapping signals, creating a fundamental bottleneck for studying cognition in ecologically valid settings. Here, we seek to isolate activation patterns during a hybrid visual and memory search task in naturalistic scenarios. We show that our deconvolution-based approach applied to coregistered EEG and eye-tracking data resolves this problem, capturing fine-grained activation patterns in the temporal response functions (TRFs) for main effects and their interactions. Starting from hypothesis-driven models, we replicated established components for visual processing and target detection in a Hybrid Search task with unrestricted eye movements. Moreover, extending our approach to hierarchically larger data-driven models enabled us to explore interactions between the effects that have otherwise been studied separately. We showed that the TRF estimates remained stable with increasing model complexity, supported by improved model performance (Pearson’s correlation coefficient) and controlled by the variance inflation factor (VIF). We identified a late activation consistent with the P300 component for target detection, and revealed that missed detections elicited similar but weaker responses, suggesting a more nuanced role than simple detection. These findings demonstrate how deconvolution methods, complemented with robust measures of model performance that support its expansion in features’ space, can uncover the dynamic interplay of attention and memory processes underlying free-viewing behavior.

## 1. Introduction

Visual search, the task of locating a target among distractors, is a well-studied paradigm in cognitive psychology, offering valuable insights into attention mechanisms. In this context, *distractors* are visual stimuli that are present in the search display but are not the goal of the search. These items compete for visual attention, challenging the participant’s ability to quickly and accurately identify the specific item they are searching for. A key manipulation in these studies involves varying the *search set size* (Wolfe, 2007), which refers to the total number of items displayed simultaneously during a visual search task. By doing so, researchers can investigate how the complexity of the visual environment impacts attention allocation and search performance. While visual search is fundamental to daily activities, it rarely occurs without the presence of multiple confounds in real-life situations. This poses the need to shift toward more ecologically (Tatler et al., 2011) valid approaches to studying visual search. For instance, while searching for a lost item closely aligns with traditional visual search experiments, many other real-world searches do not fit this paradigm. Searching for products in a grocery store, selecting clothes in a shop, or looking for one of several books of interest in a bookstore each involves distinct cognitive demands. Similarly, finding one of your friends in a crowded place requires integrating contextual cues, memory, and decision-making in ways that go far beyond the simplified conditions of laboratory tasks. When many possible targets must be held in working memory, the task is called Hybrid Search (HS), because the search must be done both in memory and in the visual scene, matching fixated items with those memorised in the first place. Thus, understanding more realistic scenarios requires considering additional factors like memory load (Schneider & Shiffrin, 1977). In HS, memory and attention are intertwined, requiring individuals to search for targets stored in memory while processing the ongoing visual information. Studies on hybrid search have expanded traditional visual search paradigms by examining how memory influences search efficiency and accuracy across varying set sizes (Barbosa et al., 2024; Wolfe, 2012).

One established way to study cognitive processes in free viewing studies has been through fixation-related potentials (FRPs), where the EEG signal is aligned to the onset of fixations and then averaged (Auerbach-Asch et al., 2020; Devillez et al., 2015; Dimigen et al., 2011; Hiebel et al., 2018; Kamienkowski et al., 2012, 2018; Kaunitz et al., 2014a; Ries et al., 2016, 2018). Analyses of dual-task paradigms have found that working memory load affects attentional control and the resolution of response conflict by diminishing the ability to ignore irrelevant distractors (Pratt et al., 2011). However, much remains to be understood about the neural dynamics underlying free-viewing tasks, particularly in complex tasks such as hybrid search. Deconvolution allows for disentangling overlapping neural activations from different sources—whether they arise from multiple stimuli or multiple cognitive processes—resulting in clearer insights into the timing and nature of the underlying brain activity (Crosse et al., 2016; Dimigen & Ehinger, 2021; Kristensen et al., 2017; N. J. Smith & Kutas, 2015b). Previous studies (Auerbach-Asch et al., 2020; Care et al., 2023; Coco et al., 2020) have demonstrated the utility of deconvolution in visual tasks including visual search, exploration and reading (Ehinger & Dimigen, 2019). For instance, Auerbach-Asch, et al. (2020) showed that the N170 face-effect is evident for the first fixation on a stimulus in free-viewing, disentangled from subsequent fixations; while Coco, et al. (2020) showed a fixation-related difference between objects’ consistency and scene context. More broadly, similar methods have been applied in the speech domain to estimate impulse responses or Temporal Response Functions (TRFs), where stimulus features represent aspects of the speech signal (Crosse et al., 2016, 2021; Desai et al., 2021; Di Liberto et al., 2015; Gonzalez et al., 2024). For instance, Di Liberto, et al. (2015) showed that combining both low-level spectrotemporal information and phonetic features best describes the relationship between neural activity and continuous speech. Desai, et al. (2021) later generalised these findings to more naturalistic stimuli. Together, these findings show the flexibility of deconvolution approaches for modelling neural activity in complex settings.

Despite these advances, deconvolution approaches still face several challenges, including issues with regularisation, goodness of fit, multi-collinearity, and interpretability (Crosse et al., 2021; Dimigen & Ehinger, 2021; Frömer et al., 2024; O’Connell et al., 2025; N. J. Smith & Kutas, 2015a, 2015b). Critically, previous efforts using a deconvolution approach expanded model complexity without controlling for multi-collinearity. As new features are incorporated into the model or the time delays for responses are increased, the number of predictors in the design matrix increases proportionally, potentially introducing collinearity problems. While measures like the variance inflation factor (VIF) have been suggested to assess collinearity in these models (N. J. Smith & Kutas, 2015b), they are rarely implemented in practice. Instead of explicitly measuring collinearity, researchers in domains such as speech processing typically address these complexities by adding a regularisation term to the regression’s loss function, penalising the addition or magnitude of new terms (Hastie et al., 2009). In particular, they use Ridge regression to return smooth temporal response functions (TRFs) (Crosse et al., 2016, 2021; Desai et al., 2021; Di Liberto et al., 2015; Gonzalez et al., 2024). This regularisation term is also employed to prevent the model from overfitting noise (Hastie et al., 2009).

TRF approaches typically focus on continuous variables, such as the envelope of speech or the spectrogram, and evaluate the models one at a time (Crosse et al., 2021). Only in some cases, more complex models are used to study the information shared by two variables (Desai et al., 2021), but only at the level of performance (r-squared). On the other hand, deconvolution methods inspired by linear models have introduced interactions between variables as well as non linear effects (N. J. Smith & Kutas, 2015a, 2015b). In particular, non linear effects are modelled using spline functions on the variable of interest values when predicting the EEG responses (N. J. Smith & Kutas, 2015b), expanding the representation of one variable on several predictors. This approach has been implemented in the Unfold toolbox, and it is typically used to model non linear dependencies of the EEG response on the saccade amplitude, among other effects (Ehinger & Dimigen, 2019). The hierarchical growth of a model through the inclusion of interactions and non linear predictors makes the evaluation of multicollinearity and appropriate regularisation even more relevant.

Beyond methodological development, recent studies have begun to address empirical questions using deconvolution methods in the domain of active vision (Auerbach-Asch et al., 2020; Care et al., 2023; Coco et al., 2020). Recently, we applied this same scheme to a visual search task in which fixation-related potentials (FRPs) to images of faces were estimated through deconvolution, allowing us to reveal a significant effect of the progression of the trial, operationalised as the number of fixation along the scanpath (trial progression score, TPS) (Care et al., 2023). In visual search, fixated stimuli are known to elicit early responses linked to low-level features (Ries et al., 2018). For instance, the P100 and N170 components from classical fixed-gaze experiments generalises to free-viewing paradigms, exhibiting reliable latency and topography (Auerbach-Asch et al., 2020; Kamienkowski et al., 2012; Kaunitz et al., 2014a). Regarding late-stage processing, gazing the target stimuli elicits modulations consistent with a P300 component after target presentation in fixed-gaze (Kamienkowski et al., 2012). The presence of distractors, manipulations on target probability, or target expectancy affected EEG responses at around 300 ms after participants locate a target in free-viewing tasks (Brouwer et al., 2013; Devillez et al., 2015; Hiebel et al., 2018; Kamienkowski et al., 2012, 2018; Kaunitz et al., 2014b; Ries et al., 2016). Moreover, working memory load has been shown to modulate these early responses, particularly when retained information involves a competing task (Pratt et al., 2011; Ries et al., 2016). Finally, the trial progression also modulates neural activity as we showed in our previous work (Care et al., 2023; Kamienkowski et al., 2018).

Our analysis strategy followed a two-step progression. Firstly, we employed a hypothesis-driven model, focusing on widely studied effects from the visual search and memory literature, such as the target-related P300 and memory load impacts on early components. This step served to validate our approach by confirming that canonical effects could be recovered under conditions of free viewing, thereby establishing coherence with prior fixed-gaze findings. Secondly, we adopted a data-driven extension, gradually enriching the models with additional predictors and interaction terms. This allowed us to explore potential effects that have not been systematically investigated in previous work. We hypothesised that main effects would remain robust and stable across models of increasing complexity, while interactions would uncover more nuanced spatiotemporal activations reflecting the interplay of attention and memory. Importantly, the predictive power of the models and the collinearity of the variables was monitored as complexity increased, ensuring interpretability while opening new opportunities to investigate unexplored interactions. By combining deconvolution with a hybrid search paradigm, we aim to advance the understanding of the neural dynamics underlying real-world search behavior and to demonstrate the value of this approach for disentangling overlapping cognitive processes.

## 2. Materials and Methods

### 2.1. Participants

The experiment involved participants from two different sites. One group was registered at the University of Buenos Aires (UBA) and the other at the University of Nottingham (UoN). All participants were naive to the experiment’s objectives, possessed normal or corrected-to-normal vision, and willingly provided written informed consent. The study was conducted in accordance with the principles embodied in the Declaration of Helsinki. Ethical approval was obtained from the respective Ethics Committees of each university (Protocol 284 from the Instituto de Investigaciones Médicas “Alfredo Lanari” – University of Buenos Aires, and Protocol F1317 from the University of Nottingham).

Nineteen adult participants performed the experiment at the University of Buenos Aires. Data from three participants were excluded: two due to bad reference signals, and one due to poor eye-tracking quality. Thus, data from 16 participants with a median age of 25 y.o. (range = [19, 40] y.o.) was analysed. Thirty adult participants performed the experiment at the University of Nottingham. Data from four participants were excluded: two due to poor eye tracking quality, and two due to poor EEG quality, even after filtering artifacts related to eye movements. This resulted in 26 participants with a median age of 20 y.o. (range = [18, 36] y.o.). Across both sites, the final dataset consisted of 42 participants.

### 2.2. Experiment

The main task involved memorising items presented during the memory phase (1, 2, or 4 items) and subsequently searching for one of those targets within a composite image during the visual search phase (Fig 1). Search images featured natural scene backgrounds, with participants asked to identify the presence or absence of a target. In the target-present condition, only one of the memorised items was present in the search image. Targets were present in 50% of the trials, creating a balanced experimental design.

**Figure 1.**
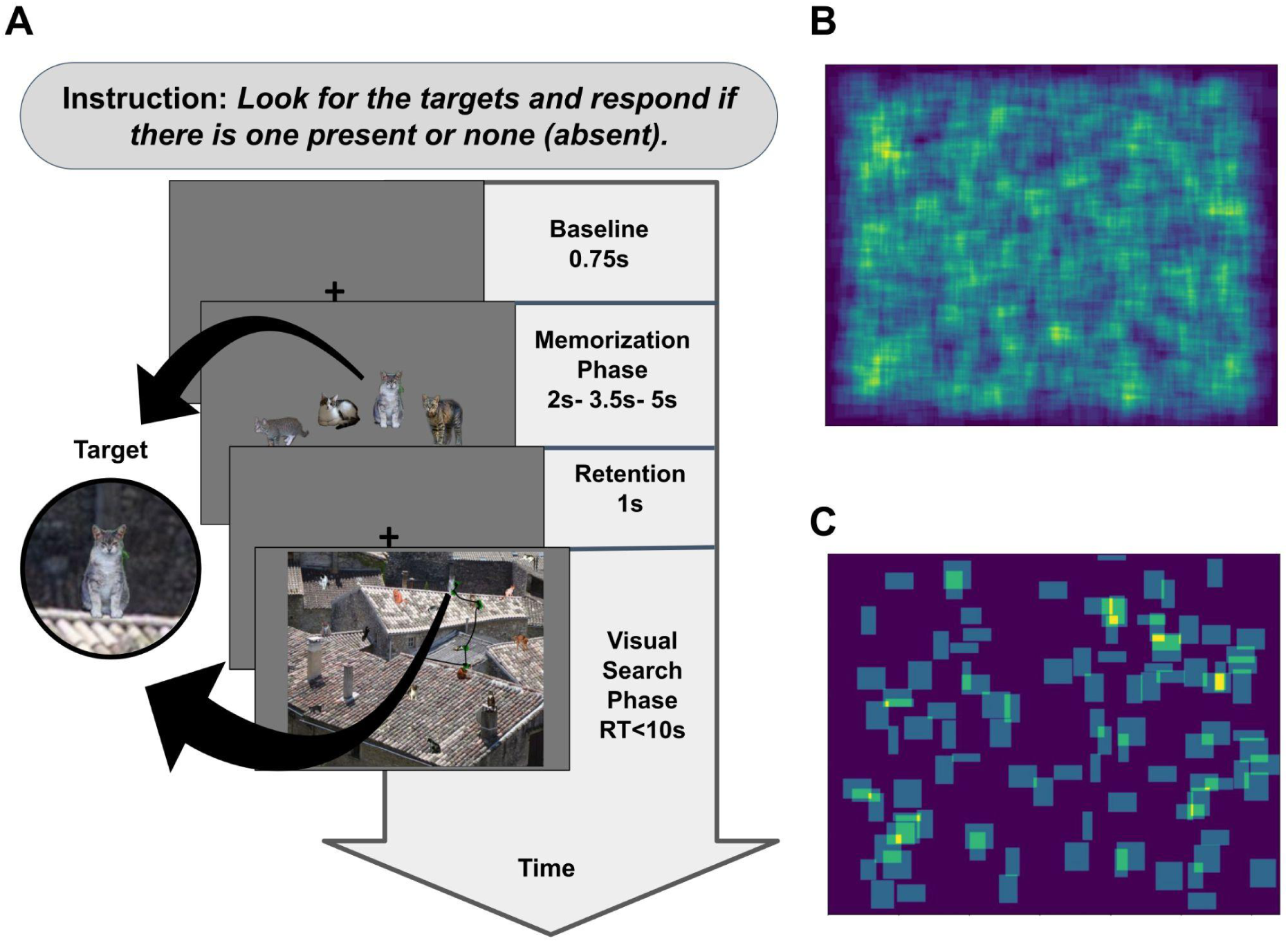
A. Experimental Paradigm. Time progression of the trial for the condition Target present (detailed in the last panel) and Memory Set Size (MSS) of four. In Target present condition only one of the potential targets was visible within the search image, while in Target absent none of them are present. In all conditions, during the Visual Search Phase, a Visual Set Size (VSS) of 16 is presented. **B-C. Bounding Boxes.** Bounding boxes for all the distractors (**B**) and the targets (**C**) presented during the Visual Search Phase across all trials and all conditions.

The trial was initiated with a fixation cross, persisting for 0.75 seconds, serving as an initial focal point (Fig. 1A). Then, there was a memorisation phase, where participants encountered different numbers of items to memorise. Both memory set sizes (MSS) and target present conditions were selected pseudo-randomly from the set {1, 2, 4} and target {present, absent}. The duration of this memorisation phase varied with memory set size, taking values of 2, 3.5, and 5 seconds for MSS values of 1, 2, and 4, respectively. After the memorisation phase, a retention period is indicated with a fixation cross for one second, and the visual search (VS) phase initiated, the visual search image persisted until participants delivered a response up to a maximum of 10 seconds.

As in previous studies (Care et al., 2023), participants completed 7 eye-map blocks throughout the experiment to characterise artifactual components. In the eye-map task, participants were instructed to blink steadily at a central point or to shift their gaze slowly between two dots displayed at fixed positions. The dots were always present during each trial at close or far positions in the vertical or horizontal axis.

The experiment was organised into 7 blocks, each comprising 30 trials, resulting in a total of 210 trials. Each visual search block was preceded by an eye-map block. The entire experimental protocol lasted approximately 50 minutes. To maintain accuracy, a drift correction was performed at the beginning of each block, opening the possibility of recalibrating if necessary. Synchronisation marks were sent from the Stimulation PC to both ET and EEG at the beginning and end of each trial, as well as at the onset of each eye-map block.

### 2.3. Stimuli

A dataset was generated, comprising 210 natural scene backgrounds with 16 items immersively integrated and semantically coherent with the contextual environment (Ruarte et al., 2025). Various categories of objects and animals were included in the dataset, utilising the annotations of the COCO dataset to acquire segmented versions of the items (Lin et al., 2014). The backgrounds were taken from ImageNet (Deng et al., 2009) dataset. Additionally, the dataset underwent a scaling process to ensure uniformity, with the larger dimension of each item set to 80 pixels, these items were also used for the presentation of potential targets (Fig. 1B,C). Following the integration of the 16 items into the backgrounds, the resulting images were standardised to dimensions of 1280 x 1024 pixels. The strategic arrangement of items was meticulously performed by hand, ensuring a sparsely distributed layout while preserving the semantic integrity of the overall image (Fig. 1).

### 2.4. Apparatus and data acquisition

The equipment used in the acquisition was the same for both datasets, except for using a cap with a different number of channels at each site. EEG activity was recorded using a BioSemi Active-Two system at 1024 Hz (64 electrode positions at UoN and 128 at UBA on a standard 10-20 montage). Four reference electrodes were used: the right and left mastoids, and the right and left earlobes. After the data were recorded, the sampling rate was digitally downsampled to 500 Hz, and the signal was re-referenced to the linked mastoids. Eye-movement tracking (ET) data were recorded using an EyeLink 1000 Plus remote system in monocular mode and a sampling rate of 500 Hz. To integrate the ET and EEG data, the stimulus-generating computer sent shared messages to both the EyeLink and BioSemi systems at different times, including the start of each trial.

At the UoN, the synchronisation was verified in a few participants using a photodiode. At the UBA, horizontal and vertical eye movements coordinates were sent analogically from the eye tracker to EEG, and added to the EEG channels before digitising them. This way, it is possible to check for delays between the EEG and the eye tracker.

In both places, the stimuli were presented in a BenQ XL2420Z monitor with a screen resolution of 1920 x 1080 pixels and at a refresh rate of 75Hz. Participants were placed at a distance of approximately 60 cm from the monitor. All stimuli were presented using Psychopy software (Peirce et al., 2019), and the responses were collected with a ‘qwerty’ keyboard.

### 2.5. EEG and eye movements pre-processing

The EEG data underwent preliminary processing using MATLAB with the EEGLAB toolbox (Delorme & Makeig, 2004) and custom scripts. Initially, a zero-phase, Hamming-windowed finite impulse response (FIR) notch filter was applied between 47.5 and 52.5 Hz using the pop_eegfiltnew function to suppress 50 Hz power line interference. The filter order was set to 846 samples, and the transition bandwidth was estimated following EEGLAB’s default specifications.

Subsequently, a zero-phase FIR band-pass filter with cutoffs at 0.1 Hz and 100 Hz was applied to remove slow drifts and high-frequency noise while preserving the spectral range of interest for event-related EEG analyses. Channels exhibiting abnormal spectra were identified as noisy and interpolated when fewer than five contiguous channels were affected. The synchronisation of eye-tracking (ET) and EEG data was accomplished using the EYE-EEG toolbox (Ehinger & Dimigen, 2019), gaze positions falling outside the screen range and blinks were flagged as erroneous data. A synchronisation quality check was performed by analysing the correlation between imported ocular data and frontal electrodes. When temporal misalignments were detected between eye-tracking and EEG signals, lag correction was implemented to ensure precise temporal correspondence between the two data streams. This correction procedure involved adjusting the relative timing of eye-tracking and EEG recordings to minimise potential time-based discrepancies that could arise from differences in data acquisition systems. Eye movements were recalculated using an offline algorithm (Engbert & Mergenthaler, 2006). Subsequently, the OPTICAT procedure (Dimigen, 2020) was applied with saccade overweighting. Following this, Independent Component Analysis (ICA) was conducted using the Infomax algorithm (Bell & Sejnowski, 1995), and Principal Component Analysis (PCA) was implemented before training in cases where data included interpolated channels.

A variance criterion (Plöchl et al., 2012) was applied to identify components associated with ocular artifacts. Manual selection of ICA projections was also performed to exclude components for which OPTICAT failed to automatically recognise ocular movement sources. The surviving ICA components were back-projected onto the EEG data, which had been high-pass filtered at 0.2 Hz and low-pass filtered at 100 Hz.

Following the initial pre-processing steps, the data underwent further preprocessing and analysis utilising Python with the MNE software (version v1.5.0) (Gramfort et al., 2013), complemented by custom functions and scripts. The 128-headcap used in UBA was mapped to the 64-headcap used in UoN for all the analyses of this work (see Supplementary Information). Fixations were categorised as landing on either distractors or targets based on a foveation criterion. Specifically, each fixation was assigned to the closest stimulus by defining a circular boundary centered on the item with a radius of 80 pixels (equivalent to 2.2 degrees of visual angle). This distance threshold was chosen to guarantee that, even if the gaze landed at the outermost edge of the boundary, some portion of the stimulus would successfully fall within the observer’s foveal region.

### 2.6. First-level Analysis: Linear Deconvolution Models for Single-Participant Data

At the individual participant analysis level, a regression-based approach was applied to estimate fixation-related responses linked to various manipulations and behavioural properties. This analytical method closely aligns with the approach detailed by (N. J. Smith & Kutas, 2015b) and the Unfold toolkit (Ehinger & Dimigen, 2019). Conceptually, it can be also viewed as a form of System Identification technique (Marmarelis, 2004), providing insights into how specific features of a stimulus map to neural responses. This method offers the additional advantage of augmenting the useful data in the analysis, thereby improving the signal-to-noise ratio by effectively handling overlapping estimations of responses.

A custom Python implementation was used to offer flexibility by accommodating different event types to specific time windows of analysis and spline modelling. This approach was instrumental in preventing the introduction of unnecessary collinearities, especially for events with shorter span time responses compared to others. The code is released and publicly available [https://github.com/incubodac/EEG_DECONV]. The variance inflation factor (VIF) method (Kutner, 1984; Marquardt, 1970) was introduced to assess the collinearity present in each model, which is important due to the increase in the number of predictors in the design matrix as function of the complexity of the model. This technique estimates collinearity by iteratively removing columns representing feature predictors and fitting models that attempt to predict the missing column using the remaining matrix. For instance, one predictor might account for a specific delay, such as estimating the response at 200 ms after a target versus a distractor appears.

Figure 2 summarizes the complete workflow of the analysis pipeline, the experimental paradigm and the corregistration to the construction of candidate models, their regularized estimation, comparative evaluation via AIC, and second level statistical analysis.

**Figure 2.**
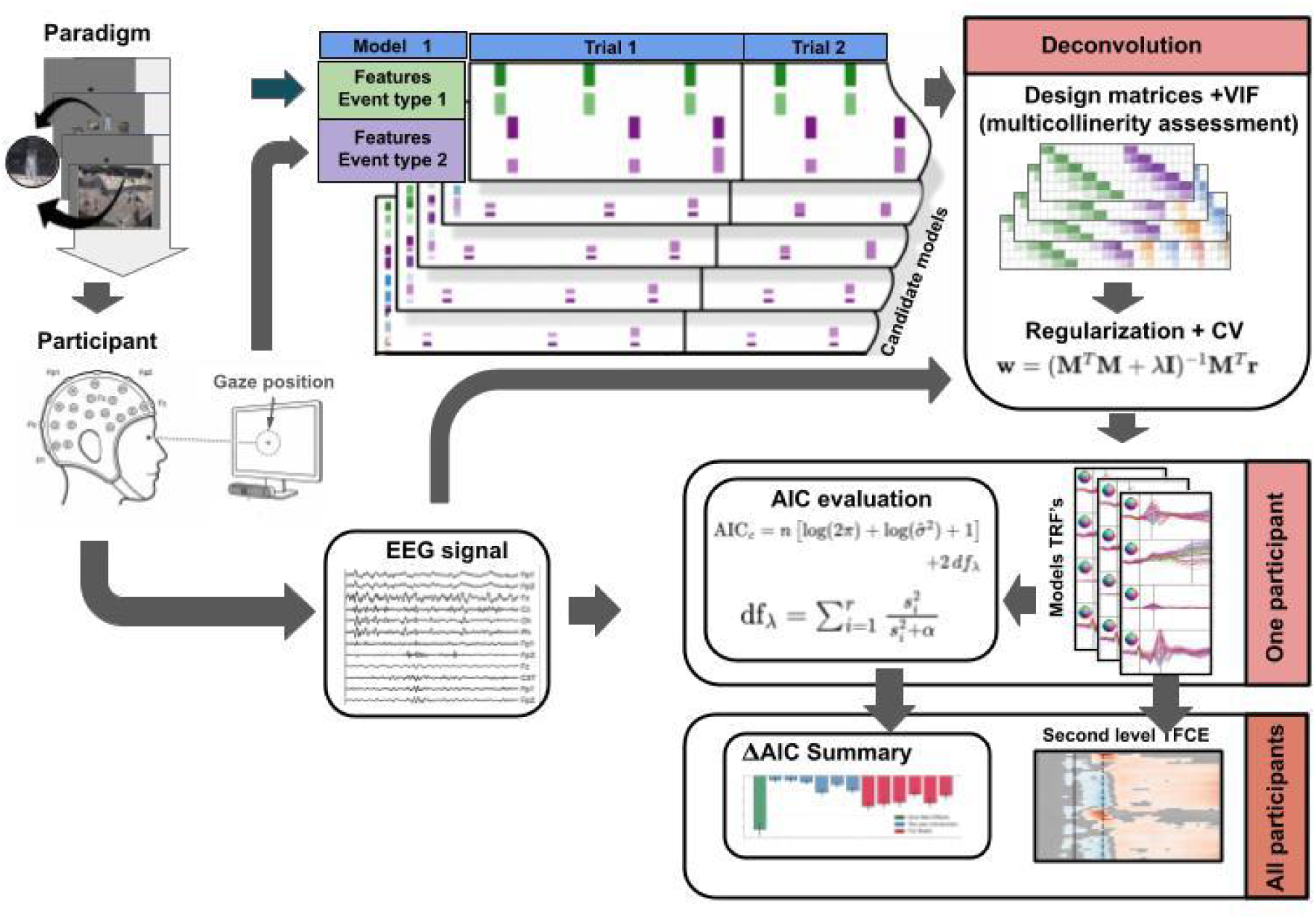
Schematic overview of the analysis pipeline. Based on the experimental paradigm manipulations and the concurrent co-registration of EEG signals and eye movement events for each participant, temporal design matrices are constructed for a range of candidate models. These model architectures are evaluated using collinearity diagnostics (VIF) and fitted via regularised deconvolution, utilizing cross-validation to select the optimal penalty parameter. The competing models are subsequently compared using Akaike Information Criteria (AIC). Finally, the estimated Temporal Response Functions (TRFs) are integrated at the group level through second-level statistical analysis.

#### 2.6.1. Models’ parameters

To explore the use of deconvolution models in this study, a regression-based approach was employed to analyse fixation-related potentials (rFRPs) in a series of models of increasing complexity. A time window of interest was defined for each participant, spanning from 200 ms before fixation onset to 400 ms afterward. Fixations classified as invalid were excluded from analysis where they landed outside the displayed image. Dummy coding was used for categorical variables like target vs distractor items, where the reference level was set to zero, allowing us to interpret parameter estimates relative to this baseline. In cases involving continuous variables, parameter estimates reflected the change in predicted neural activity measured by the EEG per unit increase in the variable.

For all the models, a saccade intercept was added to model the mean contribution of saccades to the signal and non linear contributions modeled by means of 5 cubic splines with saccade amplitude as a predictor.

The effect of the target was analysed, treating nontarget objects as the reference level. This was followed by examining effects related to the progress of the task; to do this, the fixation rank standardised per trial was used as a continuous predictor. Each participant and electrode was analysed separately using these models. Finally, baseline correction of [-200, 0] ms window was applied before visualisation and second-level analysis (see below).

#### 2.6.2. Akaike Information Criteria (AIC)

To evaluate model performance, considering the balance between goodness-of-fit with model complexity, the Akaike Information Criterion (AIC) was employed. Rooted in information theory, AIC provides an estimate of the expected relative Kullback-Leibler (K-L) distance, essentially the "information lost", between a candidate approximating model and the unknown true mechanism that generated the data. Because truth is assumed to be far more complex than any fitted model, AIC allows us to identify which model among a candidate set is estimated to be "closest" to full reality by selecting the one with the smallest AIC value (Burnham et al., 2011; Burnham & Anderson, 2004).

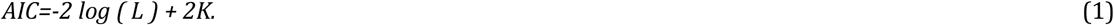

Where *L* represents the maximized likelihood of the model and *K* represents the total number of estimable parameters, including σ. The equation 1 explicitly captures the bias-variance trade-off: the first term decreases as the model fit improves, while the *2K* penalty increases with added complexity.

However, because the resultant temporal response functions were estimated using Ridge (Tikhonov) regularisation, a traditional count of estimated parameters (*K*) is inadequate and would over-penalise the model’s complexity. Instead, it is necessary to determine the effective degrees of freedom for the linear system (Zou et al., 2007). Following the trace calculation (see Supplementary Information) for Ridge regression, the effective degrees of freedom, *df(λ)* (eq. 2), can be estimated utilising the singular values (*d_i_*) of the designed matrix and the regularisation parameter (*λ*). This yields the final penalisation formula:

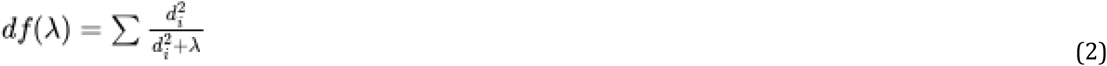

Finally, to evaluate whether incrementally adding complexity (such as new main effects or interaction terms) meaningfully improved the model, the relative AIC differences were computed between competing models, for each channel channel (*ΔAIC_ch_*). Because visual search variables impact the brain spatially, adding a specific effect (like a "Target" effect) might significantly improve the model fit in certain relevant channels while simultaneously penalising the fit in irrelevant channels. To account for this spatial variability, the *ΔAIC_ch_* was aggregated for each subject and model comparison, and selected the lowest *N* values of *ΔAIC_ch_* representing the channels that benefited the most from the added complexity. These channel-wise values were averaged for each participant, allowing us to evaluate the final mean *ΔAIC* across subjects to allow comparisons between models of different complexity.

#### 2.6.3. Variance Inflation Factor (VIF)

The Variance Inflation Factor (VIF) (eq. 3) is a measure used to assess the extent to which the variance of a regression coefficient is inflated due to multicollinearity among predictors. Crucially, collinearity directly affects the noise in the model’s estimates and dictates how much data is needed to achieve reliable results; therefore, examining VIFs is key to understanding and evaluating this impact (Smith & Kutas, 2015). Because continuous EEG noise exhibits strong temporal autocorrelations, computing mathematically exact VIFs (which assume independent errors) is currently unfeasible (Smith & Kutas, 2015). To evaluate multi-collinearity in our deconvolution model, we instead calculated and averaged independent VIFs across time lags for each predictor. This averaged estimation serves as a robust and practical proxy for diagnosing problematic collinearity. To calculate the independent VIF for a given predictor, the predictor is regressed against all other predictors in the model, and the resulting R2 value is used in the formula:

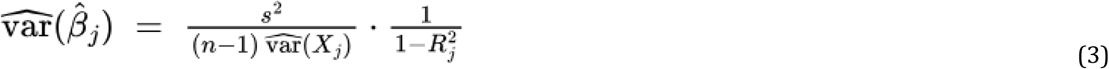

Where *R ^2^* is the coefficient of determination from this regression. A high VIF value indicates substantial multicollinearity, suggesting that the variance of the coefficient estimate is inflated due to strong correlation between a predictor and other predictors. In practice, a VIF greater than 10 is often considered indicative of problematic collinearity requiring further investigation or remediation (Kutner, 1984).

#### 2.6.4. Regularisation

L2 regularization (Ridge regression) was used to mitigate the variance inflation caused by multicollinearity. Ridge regression preserves the linear structure and interpretability of the model while a penalty on coefficient magnitude for the linear regression. This regularisation term helps to stabilise the estimates and reduce the impact of multicollinearity, leading to more reliable and generalisable models (Crosse et al., 2016; Hoerl & Kennard, 1970)(Crosse et al. 2016, Kennard 1970).

To determine the optimal L2 regularisation penalty (denoted here as α) for each specific model and participant, a cross-validation procedure was employed. Because the ideal regularisation strength depends heavily on the unique noise characteristics and the number of available trials for a given participant, this parameter must be tuned individually to optimally balance the bias-variance tradeoff. For each subject and model architecture, the data was partitioned using a 10-fold cross-validation scheme. Within each of the 10 iterations, the Ridge regression model was trained on 90% of the data across a predefined grid of α values. The predictive performance of these fitted models was then systematically evaluated on the unseen 10% of the data (the test fold). To assess this goodness-of-fit,the Pearson correlation coefficient (R) was calculated between the model’s predicted EEG responses and the actual, recorded continuous EEG signal in the test set. The α value that yielded the highest mean Pearson correlation across all 10 test folds was identified as the optimal regularisation parameter for that specific subject and model. Finally, to leverage the full power of the recordings, the model was completely retrained on the entire dataset utilising this best-performing α. The regression coefficients (TRFs) extracted from this final retrained model were the ones used for all subsequent group-level statistics and topographical visualisations.

#### 2.6.5. Predictors

##### Fixation Intercept

The fixation intercept encoded whether subjects fixated on a stimulus item by defining a circular patch centered on each item, with a radius of 80 pixels. This radius was chosen to ensure that, even at the outermost edge of the circle, some part of the stimulus would likely be within the foveal region. Fixations were classified as on-stimulus if they were contained within the circular region.

The fixation intercept predictor serves as a baseline or reference condition, representing the response when other predictors are set to zero. Unlike a traditional bias term in linear models, this intercept incorporates the deconvolved estimates of the base condition. This predictor is aligned with the fixation onset, and spans through the time-lag window used in the TRF (temporal response function) analysis.

##### Target

Fixations falling within a target’s patch were labeled as "target fixations," while those within a distractor’s patch were labeled as "distractor fixations." This classification method allowed for accurate distinction of fixation events based on the type of item each fixation targeted.

##### Standardised Rank (TPS)

The fixation rank within each trial was used as a predictor to characterise the sequence of fixations. Since trial durations varied based on each subject’s response time, the fixation rank was standardised (called trial progress score or TPS), providing a consistent measure of the processing progress within each trial (Care et al., 2023). The distribution of the TPS is similar across both datasets.

##### Memory Set Size (MSS)

The different values of the memory set size (MSS) was encoded as an ordinal predictor with levels {0, 1, 3}. This encoding ensured that the intervals between predictor values remained consistent, while shifting the base condition (MSS = 1) to the fixation intercept.

##### Saccade intercept and Amplitude

To complement fixation-related predictors, saccade-related predictors were also included. The saccade intercept captures the baseline response associated with saccadic movement with the portion of the lambda response related to the saccade (Kaunitz et al., 2014a; Thickbroom & Mastaglia, 1990), while the saccade amplitude predictor quantifies the distance covered by each saccade. Given that the relationship between saccade amplitude and electrophysiological response is known to be non linear, the contribution of saccade amplitude was modeled using five cubic B-splines. This approach provides a flexible, smooth fit to capture non linearities in the response associated with different saccade amplitudes. This spline-based approach was applied across all models (Ehinger & Dimigen, 2019; Tremblay & Newman, 2015). In brief, the spline basis was constructed using the BSpline implementation from the SciPy v1.10.1 library. A cubic spline order was used (k=3), with three internal knots placed at the empirical quantiles of the observed amplitude distribution to ensure balanced coverage across the data range. Boundary knots were placed at the minimum and maximum observed amplitudes and replicated three times at each end, as is standard for cubic B-spline basis construction, to ensure the basis spans the full data range. This configuration resulted in a basis of five spline functions. For interpretability, one spline basis function (the one with the largest value at the mean saccade amplitude) was removed from the design matrix and replaced with an intercept term, allowing the estimated coefficients to reflect deviations relative to typical saccade magnitudes.

##### Models

Our hypothesis-driven model was defined as:

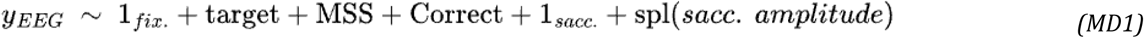

A full model was also implemented, incorporating all first-order interactions and spline expansions:

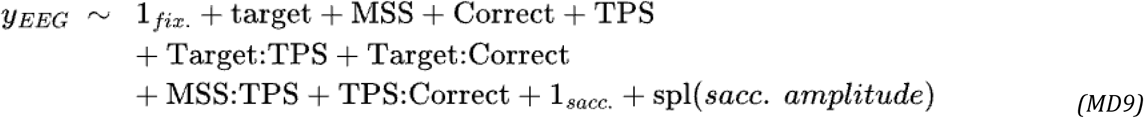

### 2.7. Second-level analysis: Statistics between participants

A second-level statistical analysis was performed using the threshold-free cluster enhancement (TFCE) (Mensen & Khatami, 2013; S. M. Smith & Nichols, 2009; Care et al., 2023). The same analysis taken in Care et al., 2023 was followed. For this analysis, the Python implementation of TFCE was used, available in MNE software (version 1.5.1) (Gramfort et al., 2013).

Figure 2 includes the comparative evaluation via AIC and second level statistical analysis in the complete workflow. In brief, this is a data-driven approach that utilises permutation-based statistics for EEG data analysis that accounts for dependencies between neighboring EEG samples in both time and space. The procedure was applied after first-level analysis, where temporal response functions (TRFs) or regression ERPs (rERPs) were obtained for each variable and participant. These estimations, which consist of the slopes (regression coefficients) for each electrode and time point, were compared against zero to test the null hypothesis (no relationship between the EEG signal and the variable of interest at each electrode and time sample).

The permutation procedure was repeated 2048 times for each cluster under the null hypothesis. The TFCE parameters were set to h = 2 (height parameter) and e = 0.5 (extent parameter), following standard recommendations in the literature (S. M. Smith & Nichols, 2009). Statistical significance was assessed using a family-wise error rate (FWER) correction at α = 0.05. It is important to point out that significant results yielded by TFCE can only support the existence of an effect related to the observed clustered data, and not its specific distribution in space and time.

## 3. Results

We first analysed participants’ behavioral and brain responses during the hybrid search task (Fig. 1) to establish the basic properties of gaze behavior and provide context for the subsequent TRF analyses.

### 3.1. Eye-movements behaviour

Eye movements exhibited typical characteristics present in free-viewing experiments (Fig. 3). The main sequence of saccades (Fig. 3A) exhibited a consistent linear relationship between peak saccade velocity and amplitude (Otero-Millan et al., 2008), and the saccade amplitude distribution showed known non linearities inherent in the saccadic system (Lebedev et al., 1996) (Fig. 3B). The distribution of fixation durations followed a characteristic positively skewed pattern pattern observed in exploration and visual search, with a peak around 160 ms and a mean duration of 219 ± 196 ms (Otero-Millan et al., 2008) (Fig. 3CD). Saccade direction displayed a preference for horizontal movements, a tendency probably accentuated by backgrounds containing a visible horizon and ground objects as distractors (Fig. 3D). The heatmap of fixations across all participants revealed widespread exploration of the visual scene (Fig. 3E). On average, participants performed 10.53 ± 2.27 fixations per trial, but the distribution of fixation ranks within trials varied systematically with memory set size (MSS), indicating that participants required progressively more fixations before responding in trials with higher memory load (Fig. 3F).

**Table 1.** Total number of trials and fixations in the whole dataset (42 participants)

| Target | Response | MSS = 1<br>(# trials / # fixs) | MSS = 2<br>(# trials / # fixs) | MSS = 4<br>(# trials / # fixs) |
| --- | --- | --- | --- | --- |
| Absent | Incorrect | 47 / 1,148 | 152 / 3,799 | 329 / 7,944 |
| Absent | Correct | 1,368 / 25,753 | 1,216 / 26,165 | 1,127 / 26,321 |
| Present | Incorrect | 244 / 3,972 | 401 / 8,657 | 649 / 15,310 |
| Present | Correct | 1,201 / 10,611 | 1,012 / 12,076 | 731 / 11,169 |

**Figure 3:**
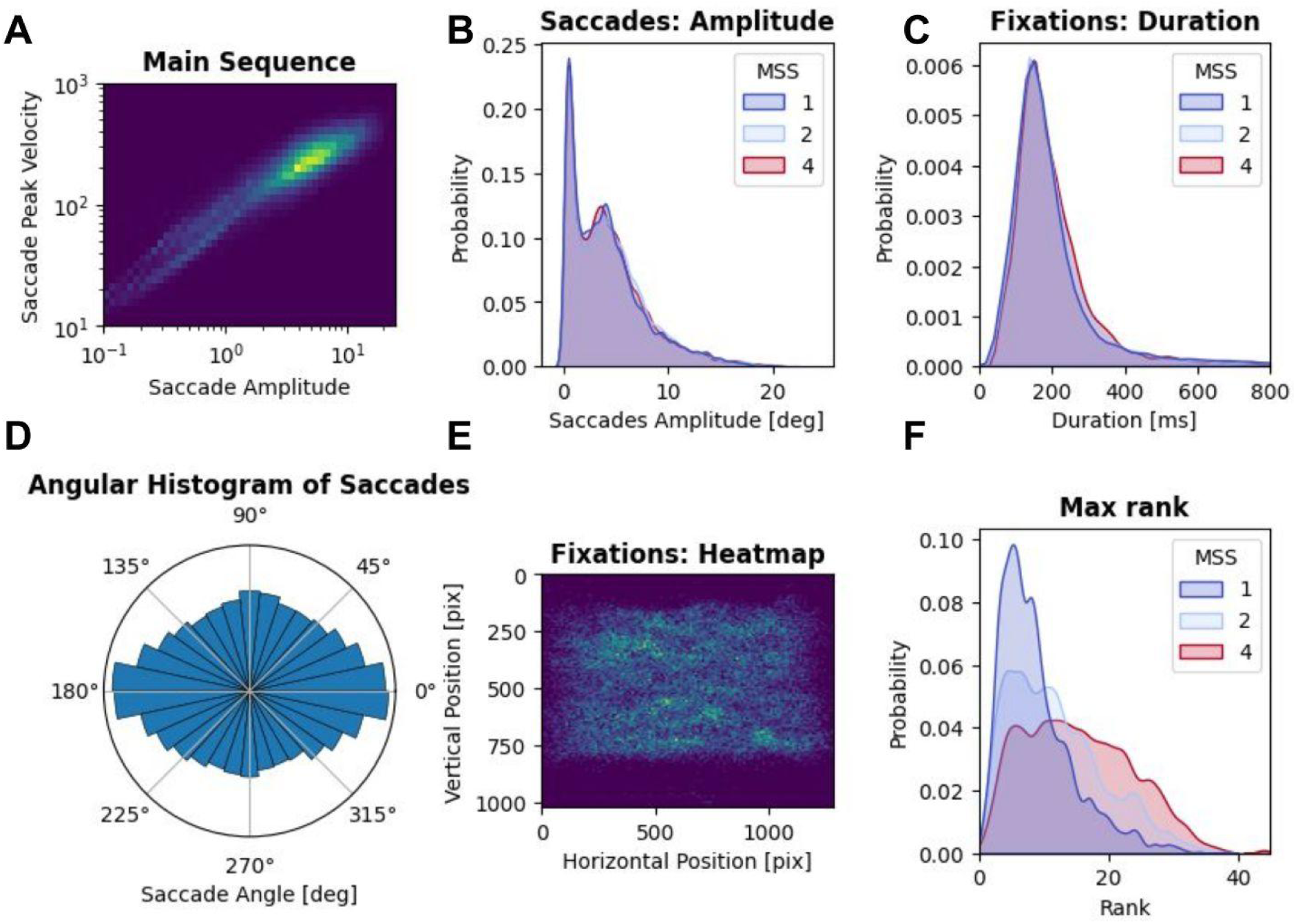
Analysis of Eye Movement Parameters During Visual Search. Top row: (A) Main sequence relationship between saccadic peak velocity and amplitude; (B) Distribution of saccade amplitudes; and (C) Fixation duration distributions, all stratified by Memory Set Size (MSS). Bottom row: (D) Angular distribution of saccadic movements; (E) Spatial distribution of fixations represented as a heatmap; and (F) Rank distribution of fixation stratified by MSS.

### 3.2. Brain Activity - Temporal Response Functions

We ran the analysis on trials in which the target was present. To investigate the activity associated with fixations and task-related constraints such as MSS, we estimated Temporal Response Functions (TRFs) using a series of models of increasing complexity. For each participant, we included variables associated with fixations such as the fixation intercept, target presence, trial progress score (TPS), and MSS; and defined a time window of interest spanning from 200 ms before to 400 ms after fixation onset. We considered the same time window aligned to the saccade onset, for the saccade intercept and the saccade amplitude. For this variable we added 5 cubic splines as predictors to cope for the non linear relation. In the present framework, the activation elicited by fixations on distractors is represented by the regression coefficients for the Intercept, while the activation elicited by fixations on target stimuli is represented by the regression coefficients for Target predictor plus the Intercept.

To test our hypotheses, we first specified a model including the predictors *Target*, *MSS*, and *Correct*, alongside the fixation and saccade intercepts, and the saccade amplitude terms (model **MD1**). Figure 4 shows the temporal activation for each coefficient and model, together with the topography of significant regions as identified by TFCE. The estimated responses confirmed that the primary effects identified are consistent with findings from other visual search experiments. We also observed the generalisation of well-established fixation-related potentials such as the P100 and the P300.

**Figure 4:**
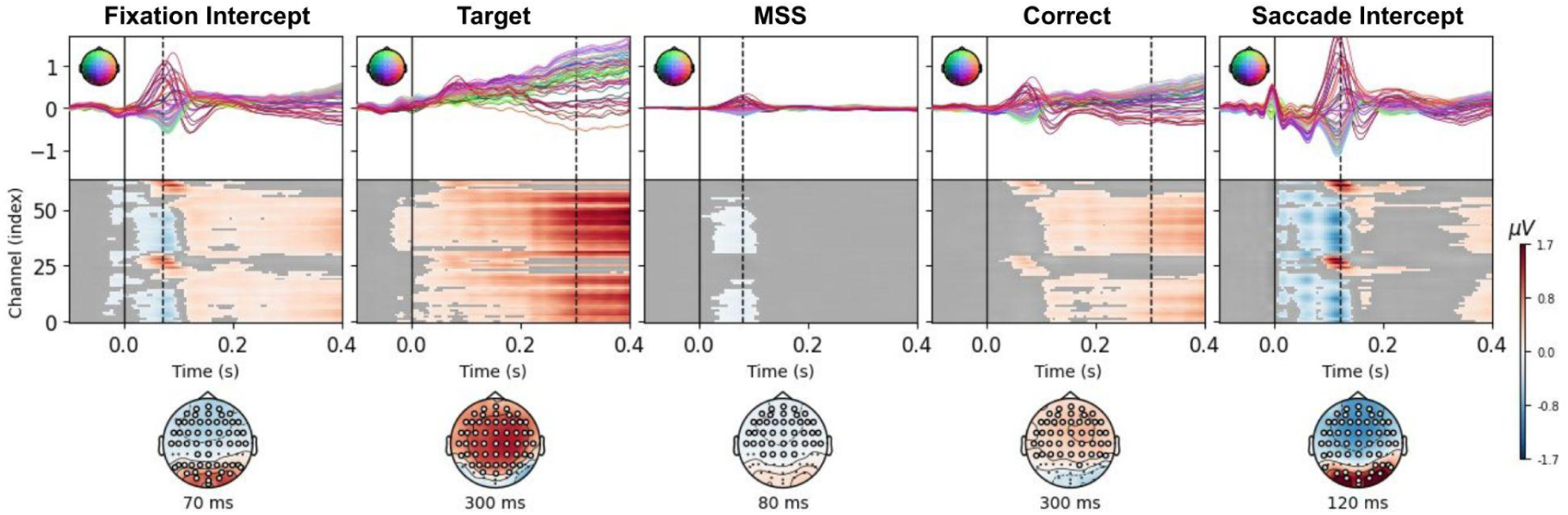
Hypothesis driven main effects and intercepts (MD1). The five effects modelled by MD1 were presented in columns: Fixations Intercept, Target, MSS, and Correct response effects aligned to fixation onsets, and the Saccade Intercept. The model was applied to all fixations from target-present trials, including incorrect trials. The top panels show the time course of activations for all electrodes, the middle panels show the activations masked by significant TFCE clusters, and the bottom panels present the spatial distributions of activations at the time points indicated by the dotted lines, with white circles highlighting electrodes that are part of significant clusters. Coding: MSS1: 0, MSS2: 1, MSS4: 3.

The fixation intercept predictor revealed an early activation peaking at approximately 80 ms, corresponding to a significant cluster with positive occipital activity and negative fronto-central activity. This timing is consistent with the P100, a well-characterised visual evoked potential. The saccade intercept showed robust activations consistent with the lambda response (Kaunitz et al., 2014a), peaking at around 120 ms. In addition, sharp transients were observed at saccade onset, closely aligning with the expected spike potential.

Target fixations (Fig. 4, second column) elicited widespread activations within the significant cluster across central, frontal, temporal, and parietal electrodes, with a pronounced P300 component, showing an increase in activity around 300 ms.

In the hybrid search (HS) task, we manipulated memory load by systematically varying the memory set size (MSS). Previous work (Pratt et al., 2011; Ries et al., 2016) has suggested that memory demands can influence early stages of information processing. Consistent with this, Fig. 4 (column 3) shows that the predictor associated with MSS modulates the amplitude of the early P100 component. The clusters in the middle row of Figure 4 display the TFCE results, with activations masked for p-values above 0.05. This model revealed a significant cluster with fronto-central activations overlapping with the P100 time window, indicating a negative modulation associated with the MSS. In previous studies (Pratt et al., 2011; Ries et al., 2016), higher memory load was associated with a decrease on the positive P100 component at occipital electrodes, In contrast, we observed a negative effect at frontal sites. Notably, a decrease at frontal sites can also be observed in Pratt et al. (2011), although this was not explored further. A significant correct-trial effect is also observed in Fig. 4 (column 4), with a topography resembling the target effect. This effect will be explored further in later sections.

Having established these primary effects with the hypothesis-driven model, we next adopted a data-driven approach by progressively increasing model complexity to explore additional effects. Model **MD1**, shown in Fig. 4, served as the baseline. We then examined the role of task progress (model **MD2**) (Care et al., 2023; Kamienkowski et al., 2018), operationalised as the standardised fixation rank per trial. This *Trial Progress Score (TPS)* represents the sequential progression of fixations within each trial. An open question is whether interactions between these predictors contribute additional explanatory power: From model **MD3** to model **MD8**, we explored the interactions between all pairs of variables. Finally, model **MD9** included all significant interactions. Unlike previous studies that introduced all possible predictors simultaneously, we implemented a stricter model evaluation procedure: controlling for model performance (R²) and variance inflation factors (VIF) (Fig. 5), and statistical significance (Fig. 6).

**Figure 5:**
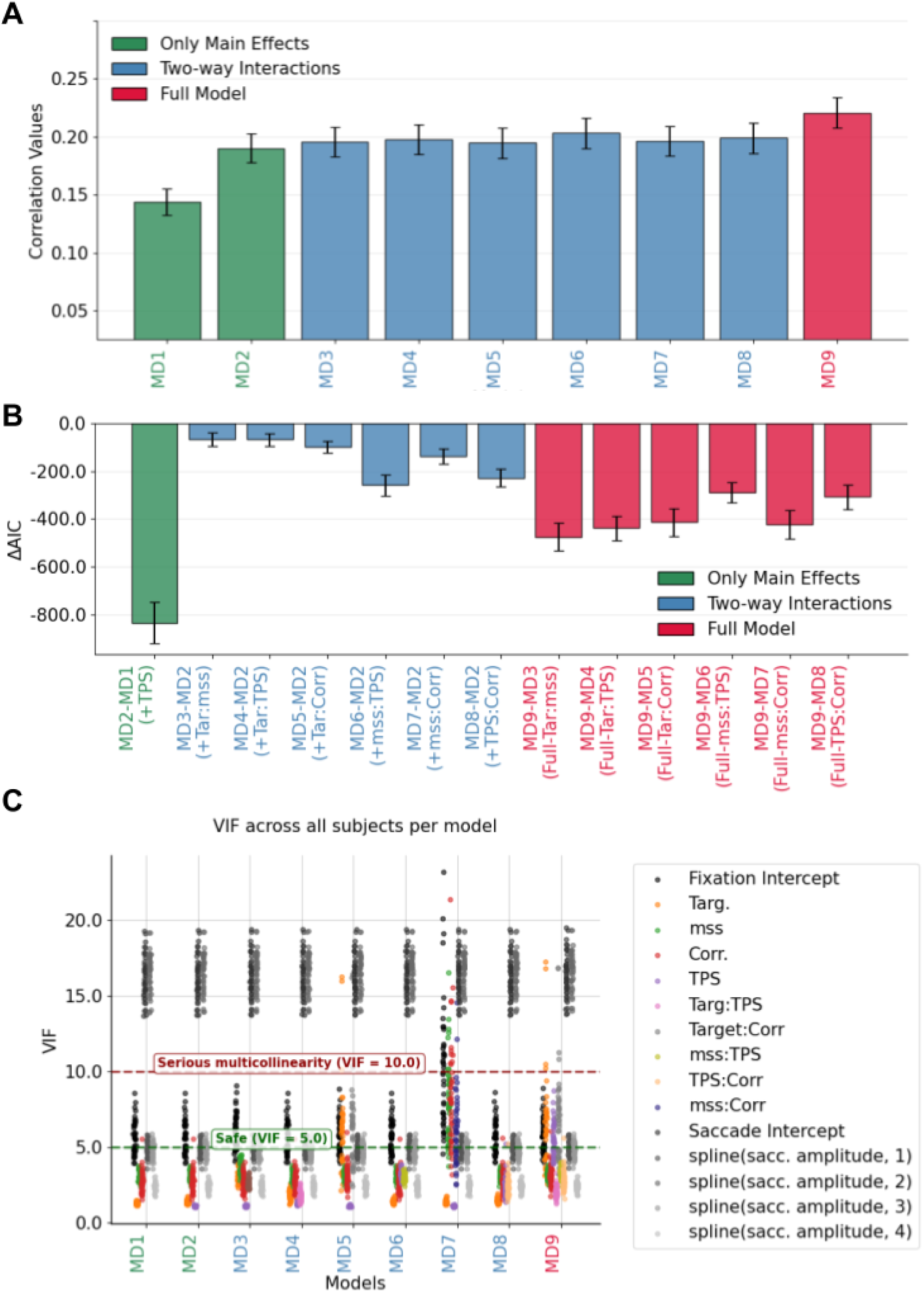
M*o*del *performance and collinearity diagnostics across subjects and model complexity.* **A.** ΔAIC between models with a different complexity was estimated for each participant and channel. Since not all of the channels are expected to respond to a given variable, only the 32 best channels were averaged per participant (similar results were obtained with all channels). Bars represent the mean (and SEM) across participants. **B.** Pearson’s correlation coefficients (*r*) were estimated, first for each channel, participant and model, and then averaged across all channels. Bars represent the mean (and SEM) across participants for each model. **C.** VIF values for each predictor, model, and subject. Distinct colors denote individual predictors, while spline-based predictors are shown in grayscale.

**Figure 6:**
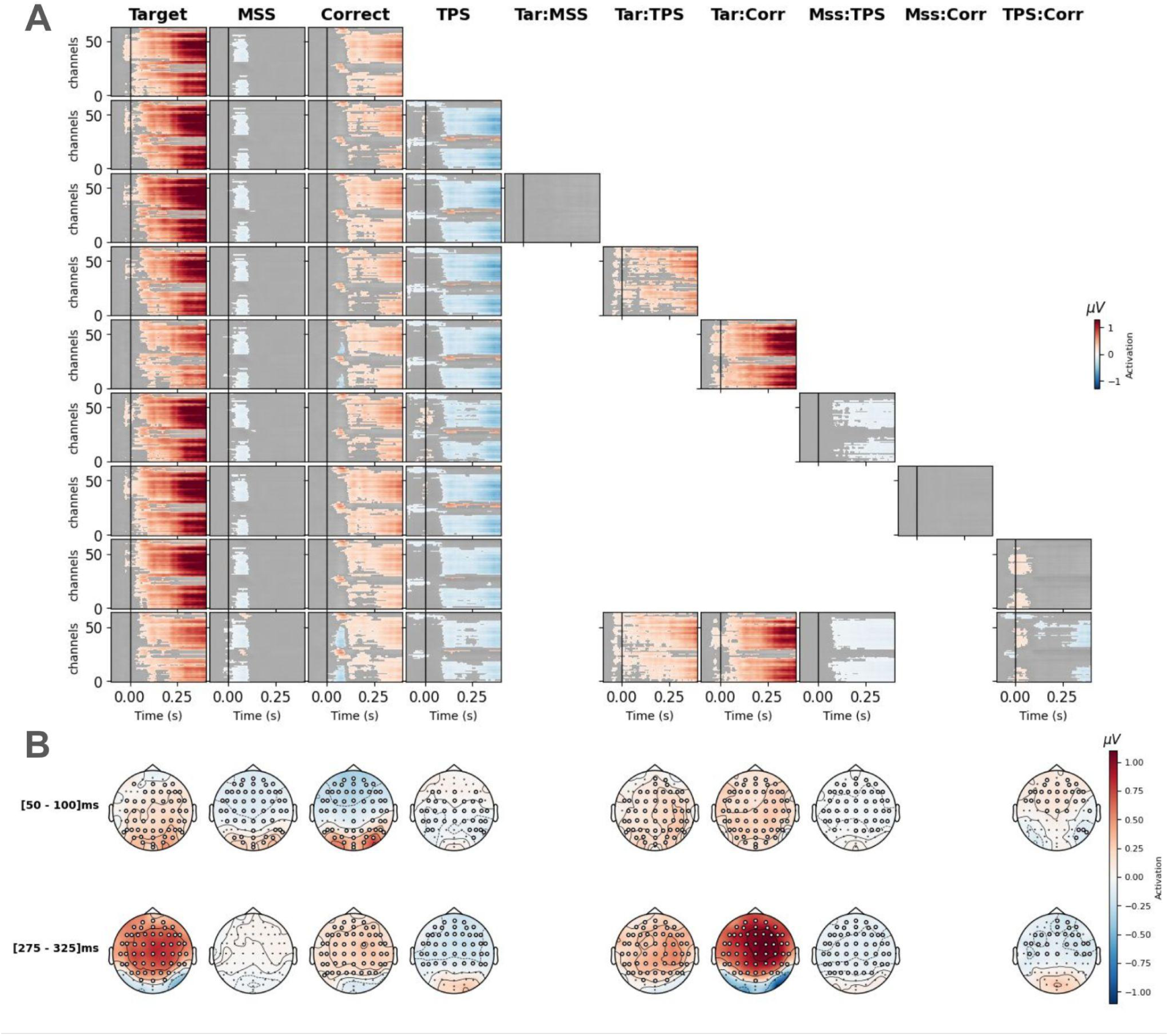
Hypothesis driven main effects and interactions (MD1 to MD9). The effects modelled in MD1-9 were presented in columns: Target, MSS, Correct response, and TPS effects, and their interactions were aligned to fixation onsets. Fixation and Saccade Intercepts were not presented here but kept the same profile as Fig. 4. The models were applied on all fixations from all target present trials, including Incorrect trials. **A.** Top panels show the activations masked by significant TFCE clusters, each row is a different model (MD1-9). **B.** Bottom panels present the spatial distributions of activations at an early ([50, 100] ms) and late ([275, 325] ms) time windows. White circles highlight electrodes that are part of the significant cluster.

Figure 5A,B shows the distribution of the Pearson correlation values across nine models of increasing complexity. In this context, the correlation is employed as a descriptive measure of how well the predicted responses match the actual EEG signal, rather than as a predictive metric on unseen data. After selecting the regularisation hyperparameter via cross-validation, the full dataset was used to compute the Pearson correlation coefficient (R) as an index of overall model fit. As expected, the simplest model (**MD1**) explains the least variance, whereas progressively more complex models capture a higher proportion of the signal, with the model including all the interactions (**MD9**) achieving the highest R values. Although the absolute correlation values are relatively low, they fall within the range typically reported in models estimating temporal response functions (TRFs) in continuous sensory perception (e.g., (Crosse et al., 2016)). This analysis demonstrates that increasing model complexity systematically improves the ability to account for EEG variance. Crucially, the concurrent application of optimal regularization ensures that this added complexity captures meaningful neural dynamics rather than overfitting to collinear noise.

Figure 5C shows the Variance Inflation Factor (VIF) for each model, predictor, and participant. For each participant, VIF values were averaged across latencies for each predictor. As expected, larger models tend to produce higher VIF values for some predictors; however, these values remain below critical thresholds in most cases. An exception occurs for predictors modeling saccades, particularly splines fit to values close to the mean saccade amplitude, which exhibit elevated collinearity. These observations further support the necessity of regularisation to stabilise parameter estimates in the presence of correlated predictors.

Figure 6 presents the estimated activations for each model (rows) and predictor (columns), with activations masked based on TFCE significance.

Including task progression (TPS predictor, model **MD2**), we observed a significant cluster with an estimated response over the occipital region, consistent across the fixation window from 70 ms to 400 ms and in line with our previous work (Care et al., 2023). Importantly, the inclusion of task progression did not substantially alter the intercepts or the main effects associated with Target fixations, MSS, and Correct responses which remained stable and robust. The figure shows activations for all channels and time points of the rERPs along with the TFCE.

Next, we explored the interactions between Target with MSS (**MD3**), TPS (**MD4**), and Correct responses (**MD5**). The activation of the interaction between Target and MSS (**MD3**) did not reach significance across participants. Conversely, the interaction between the Target and TPS (**MD4**) revealed a significant effect at frontal electrodes, with a gradual increase in activation across the fixation period. Previous effects related to intercepts, targets, and MSS remained stable, with no substantial changes. The interaction between the Target and Correct (**MD5**) also showed a significant effect with a gradual increase in activation across the fixation period. Notably, the significant effects already identified in the simpler models are consistently preserved when moving to larger models, indicating the robustness of the findings. At the same time, we observed that the interactions *Target:MSS* (**MD3**) and *MSS:Correct* (**MD7**) did not reach significance and therefore were not retained in the final and largest model. Conversely, *Target:TPS* (**MD4**) and *Target:Correct* (**MD5**) showed broad significant clusters, both characterised by gradually increasing activations during the fixation period, with the latter effect being slightly stronger. Although this interaction shows a strong statistical significance, the overall model (Fig. 5A,B) does not exhibit a substantial improvement, as it accounts for an effect that was already partially captured by previous models. The *MSS:TPS* interaction (**MD6**) yielded significant negative activations over frontal electrodes, while *TPS:Correct* (**MD8**) produced a mild but consistent positive activation restricted to occipital regions.

Finally, the full model (**MD9**), which included all significant effects, reproduced the same activation patterns observed in the preceding models, and revealed a late negative frontal modulation associated with the *TPS:Correct* interaction.

Interestingly, when adding the *Target:Correct* interaction (**MD5**, **MD9**) the main effect of *Target* is reduced and the *Target:Correct* interaction presents a strong late fronto-central positive effect (Fig. 6B). This indicates that the larger contribution to the effect is from the Correct trials. To reconstruct the Target effect on correct trials we add both *Target* and *Target:Correct* effects (Fig. 7), while the *Target* effect alone in **MD9** represents the response to targets on incorrect trials. The reconstructed effect for targets on correct trials presents a classical P300 component, while the reconstructed effect for targets on incorrect trials also presents a significant, although mild, response, consistent with previous results for missed targets on free eye movement visual search (Dias et al., 2013).

**Figure 7:**
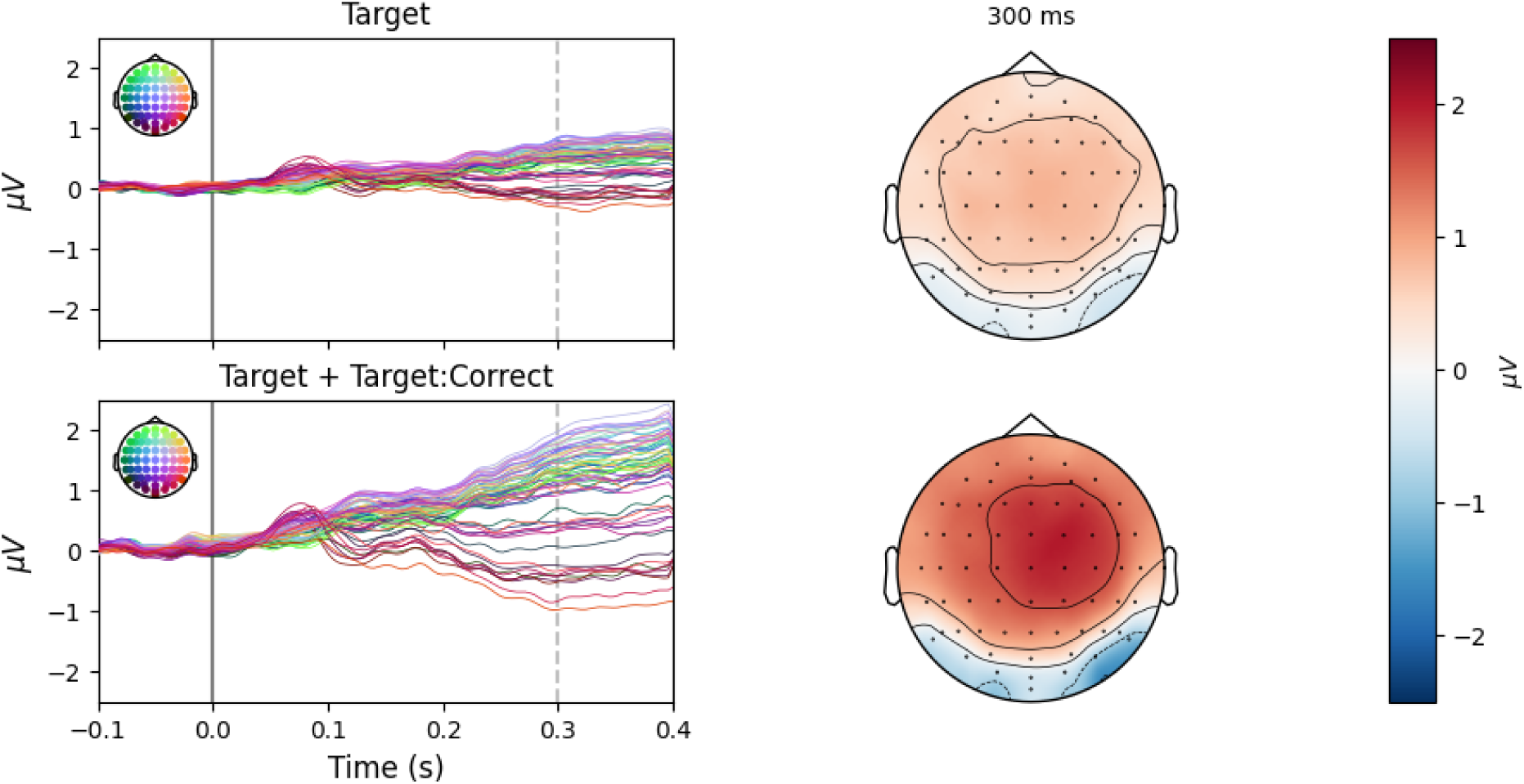
Reconstructed responses for Targets. The responses were reconstructed from the M9 model regression coefficients. The effect on Correct trials corresponds to the addition of the effects of Target==1 and Target*Correct==1. Conversely, the effect on Incorrect trials corresponds to the addition of the effect of Target==1 alone (i.e. Target*Correct==0). **A**,**B.** time course of reconstructed responses for targets on incorrect **(A)** and correct **(B)** trials. **C**,**D.** spatial distributions of reconstructed responses for targets on incorrect **(C)** and correct **(D)** trials, at 350 ms.

## 4. Discussion

This study investigated the neural dynamics of hybrid search by combining concurrent EEG and eye-tracking with deconvolution modeling, yielding significant theoretical and methodological contributions. Methodologically, it demonstrates how regularised linear models can successfully disentangle overlapping neural responses in free-viewing paradigms, while theoretically, it shows a previously unknown interplay between memory load and visual attention.

From the hypothesis-driven model (**MD1**), the analysis of the Target and Correct conditions successfully replicated established target-detection effects, such as the P300 (Brouwer et al., 2013; Devillez et al., 2015; Hiebel et al., 2018; Kamienkowski et al., 2012, 2018; Kaunitz et al., 2014b; Ries et al., 2016). However, the most novel effect relates to the Memory Set Size (MSS), which could be integrated into Lavie’s attention load theory. A key question is whether memory set size manipulation acts as a competing perceptual resource or an internally driven control demand (Lavie, 2010). Our findings show that early fixation-related activity increases with the MSS; within Lavie’s framework, this reflects increased cognitive control rather than reduced perceptual capacity. In memory-guided search, larger memory sets amplify the number of potential template mismatches and distractor pressure. Consequently, the fronto-central MSS modulation observed here likely reflects enhanced top-down control required to maintain task-relevant representations during the evaluation of each fixation. At first sight, these findings seem to contrast with previous studies exploring effects of working memory load on early visual responses (Pratt et al., 2011; Ries et al., 2016). However, in those studies, memory load was manipulated using a secondary task, creating an external resource competition that diverted processing capacity away from the primary visual task. Conversely, in the present hybrid search, memory load represents an internally generated task demand necessary to perform the primary search. This distinction between external vs internal resources’ demands provides a plausible explanation for the opposite effects of increased cognitive load found in the literature (Pratt et al., 2011; Ries et al., 2016).

Recent magnetoencephalography (MEG) research on a free-viewing hybrid search paradigm with the same stimuli provides complementary information on these early effects (Gonzalez et al., 2025). They found that, at approximately 100 ms post-fixation, frontoparietal beta-band activity scales significantly with memory load, coinciding in latency with the early visually evoked responses observed here. Traditional theoretical frameworks propose that early visual components act as a top-down filter to silence irrelevant inputs and increase the signal-to-noise ratio (SNR) for task-relevant features, a mechanism originally linked to alpha-band dynamics in fixed-gaze tasks (Klimesch, 2011). However, the concurrent MEG data suggests that during active, naturalistic search, this same cognitive control mechanism is orchestrated by beta-band networks. Therefore, the early memory load modulation observed in our fixation-related potentials likely reflects this beta-driven top-down control, dynamically increasing the SNR at the very beginning of each fixation to successfully evaluate complex sensory inputs against representations held in working memory.

In this study we also explored data-driven models, by expanding the initial model (**MD1**) with new main effects and interactions. The analysis first revealed broad effects linked to the number of fixations that occurred during a trial This effect was seen on fronto-central electrodes, coinciding with our previous findings in visual search (Care et al., 2023). The dependence of brain activity with the progression of the task can be interpreted as continuous updates to predictions regarding the potential target location and the ongoing evaluation of target presence (Care et al., 2023). Moreover, consistent with the specific demands of the current experiment, which requires observers to continuously match incoming visual input against a working memory set, this progressive evaluation and updating process was larger in higher MSS trials, as reflected in the MSS × TPS interaction. The interaction term Target × TPS suggests that target evaluation is not stationary but continuously modulated by task progression, showing gradually increasing frontal activations as trials unfold. This is consistent with increasing expectancy, as shown before in visual search tasks (Care et al., 2023; Kamienkowski et al., 2018). For instance, in oddball experiments, late components such as the P3, have been related to the concept of expectancy, as the participant is anticipating a target to be presented, a build-up of expectancy is sought to occur (Care et al., 2023; Kamienkowski et al., 2018; Polich, 2007). Furthermore, TPS showed significant interaction with the Correct trial effect, indicating a relation between trial progression and behavioral outcomes. Finally, including the Target × Correct interaction term clearly separates different contributions to the overall effect of fixating the target. This reveals that successful target detection drives the majority of the neural activity, while successfully isolating the distinct, weaker brain response to targets that were fixated but not reported (missed) (Dias et al., 2013).

Methodologically, the regression framework reveals that effects previously treated separately in classical analyses emerge naturally when modeled jointly. However, this flexibility introduces additional methodological considerations: without proper goodness-of-fit assessment, regression-based models risk overfitting, yield unstable coefficient estimates, or obscure neural effects behind collinear predictors (pitfalls that are less salient in classical univariate approaches). Moreover, we suggest that model quality requires looking beyond explained variance (*R^2^*) or correlation (*r*). This can be done by incorporating Variance Inflation Factors (VIF) to evaluate multi-collinearity before assessing model’s performance, as originally suggested by Smith and Kutas (2015). Strikingly, this has not, to our knowledge, yet been implemented in the field. VIF is particularly important as models grow in complexity, from a few predictors to larger models including non linear effects such as splines. By quantifying redundancy among predictors, this methodological tool ensures that the predictors remain stable and interpretable. In addition, using unbiased measures for model selection such as AIC is more adequate to compare growing models than using only correlation values. This brings additional challenges since using Ridge regression reduces the degrees of freedom, making its estimation non-trivial (Hastie et al., 2009; Zou et al., 2007). Here, we included all these measures, using VIF to assess multicollinearity across increasingly complex models, comparing AIC values for model selection, and estimated the optimal values for ridge regression based on the reduced number of degrees of freedom. This was optimised to run without the need of a specialised high performance server (implemented in a standard desktop workstation AMD Ryzen 5 5600X, RAM 64 Gb, NVIDIA GeForce RTX 3060 12Gb).

Although we presented several improvements to allow the analyses of more naturalistic scenarios, this approach still has some limitations that must be acknowledged. First, using natural stimuli comes at the price of potential imbalances across conditions. This can result in some conditions being over-represented while others remain sparsely sampled, posing challenges for statistical power. Moreover, allowing free eye movements throughout the scene implies that each event is unique, landing on a specific location in relation to the objects in the scene, for a specific time, and after a particular scanpath. This prevents noise ceiling techniques based on trial repetition used to reduce the variability of the signal and improve model evaluation (Chalehchaleh et al., 2025). Additionally, the findings are restricted by the spatial resolution of EEG and its limited sensitivity to higher frequency activity. These limitations could, in principle, be mitigated by using high density MEG, which may allow researchers to disentangle processes that can not be separated in the time-domain. Future research could also combine frequency-domain analyses with deconvolution to explore oscillatory dynamics more thoroughly (Litvak et al., 2013).

In conclusion, this study offers important theoretical insights into hybrid search while making a methodological contribution, by demonstrating how effects traditionally studied in isolation under fixed-gaze emerge naturally when modeled jointly. This integrative modeling framework is capable of parsing complex, overlapping neural operations inherent to naturalistic visual search.

## Acknowledgements

J.E.K received research grants from CONICET (PIP 11220150100787CO) and ANPCyT (PICT 2018-2699). J.E.K. and M.J.I. received an award from ARL (Cooperative Agreement Number W911NF2120237). We thank Alessandra Barbosa for their collaboration with the data acquisition, and Juan Octavio Castro, Gonzalo Ruarte and Anthony Ries for insightful discussions.

## Ethical statement

The study was conducted in accordance with the principles embodied in the Declaration of Helsinki. Ethical approval was obtained from the respective local Ethics Committees at each institution (Protocol 284 from the Instituto de Investigaciones Médicas “Alfredo Lanari” – University of Buenos Aires, and Protocol F1317, School of Psychology Ethics Panel, University of Nottingham). All participants provided written informed consent prior to participation.

