## Supplementary Information for "Time–space signatures of hybrid search resolution using EEG and eye movements concurrent recordings"

### Supplemental information

#### Spatial Downsampling for Cross-Montage Comparison

Since different caps were used at the two recording locations, to enable seamless comparison across different electrode layouts and conduct the final analysis on a 64-electrode montage, a bespoke function was crafted. This function is tailored to generate an interpolated version of a 64-electrode montage, using the available 128-electrode dataset. We preserved spatially corresponding electrodes between the two montages while spherical splines interpolation (Perrin et al., 1989) was used to estimate the field at the new location.

To achieve this, a virtual montage comprising 158 channels was devised. This involved incorporating the locations of the 30 non-matching electrodes from the 64 Biosemi montage into the 128-electrode montage used for recording. Subsequently, interpolation was applied to fill in the missing data, and the non-matching locations were carefully removed, resulting in the 64-electrode montage for the subsequent analysis.

#### AIC Estimation for Ridge Regression

This section derives the Akaike Information Criterion (AIC) for models fitted using Tikhonov regularization (ridge regression). The key challenge is that ridge regression does not have a well-defined integer number of parameters; we therefore replace the parameter count with the effective degrees of freedom, estimated from the hat matrix of the regularized fit.

#### 1. Effective Degrees of Freedom for Ridge Regression

Following Zou, Hastie, and Tibshirani (2007), for a fitting method that produces fitted values $\hat{y}$ from observations $y \sim N(\mu, \sigma^2 I)$, the effective degrees of freedom are defined as:

$$df = (1/\sigma^2)\ \Sigma_i \ Cov(\hat{y}_i, y_i)$$

For ordinary linear regression, $\hat{y} = Hy$ where $H = X(X^T X)^{-1} X^T$ is the hat matrix and X is independent of $\varepsilon$. Then:

$$Cov(\hat{y}, y) = Cov(Hy, y) = H\ Cov(y, y) = \sigma^2 H$$

so that $df = tr(H)$. For ridge regression with penalty parameter $\lambda \geq 0$, the hat umatrix is:

$$H\_\lambda = X(X^T X + \lambda I)^{-1} X^T$$

and therefore the effective degrees of freedom are:

$$df(\lambda) = tr(H\_\lambda) = tr[X(X^T X + \lambda I)^{-1} X^T] \quad (1)$$

#### 2. Trace via the Singular Value Decomposition

Let $X = UDV^T$ be the singular value decomposition (SVD) of X, where U and V are orthogonal matrices and $D = diag(d_1, \ldots, d_{\square})$ contains the singular values. Then:

$$X^T X = VD^2 V^T$$

$$X^T X + \lambda I = V(D^2 + \lambda I)V^T$$

$$(X^T X + \lambda I)^{-1} = V(D^2 + \lambda I)^{-1} V^T$$

$$H\_\lambda = X(X^T X + \lambda I)^{-1} X^T = UDV^T \cdot V(D^2 + \lambda I)^{-1} V^T \cdot VDU^T = UD(D^2 + \lambda I)^{-1} DU^T$$

Taking the trace and using the cyclic property and orthonormality of U:

$$tr(H\_\lambda) = tr[D^2(D^2 + \lambda I)^{-1}] = \Sigma_{\square}\ d_{\square}^2/(d_{\square}^2 + \lambda)$$

Substituting back into Eq. (1):

$$\mathrm{df}(\lambda) = \Sigma_{□}\ d_{□}^2/(d_{□}^2 + \lambda) \quad (2)$$

where the sum runs over all singular values $d_{□}$ of X. When $\lambda = 0$ this reduces to df = rank(X) = p, recovering the ordinary least-squares result.

### 3. Akaike Information Criterion

The AIC is defined as (Burnham & Anderson, 2002):

$$\mathrm{AIC} = -2 \log L(\hat{\theta}|y) + 2k \quad (3)$$

where $\log L(\hat{\theta}|y)$ is the maximized log-likelihood and k is the number of free parameters. Adapting this for ridge regression, following the proposal of Zou et al. (2007) for the LASSO, we replace the integer parameter count k with the effective degrees of freedom from Eq. (2).

### 4. Channel-wise AIC with Estimated Variance

Consider a multivariate response with C channels and n observations. We relax the assumption of a common noise variance across channels, and instead estimate $\sigma^2\_c$ independently for each channel c. The log-likelihood for channel c under a Gaussian model is:

$$\log L\_c = -(n/2) \log(2\pi\sigma^2\_c) - RSS\_c/(2\sigma^2\_c)$$

where $RSS\_c = \|y\_c - \hat{y}\_c\|^2$ is the residual sum of squares for channel c. Maximizing over $\sigma^2\_c$ yields the maximum-likelihood estimate $\hat{\sigma}^2\_c = RSS\_c / n$, giving the profile log-likelihood:

$$\log L\_c(\hat{\sigma}^2\_c) = -(n/2)[1 + \log(2\pi) + \log(RSS\_c/n)]$$

Applying Eq. (3) and counting $\sigma^2\_c$ as an additional estimated parameter, the AIC for channel c is:

$$AIC\_c = -2 \log L\_c(\hat{\sigma}^2\_c) + 2[\mathrm{df}(\lambda) + 1]$$

Expanding:

$$AIC\_c = n \log(RSS\_c/n) + n[1 + \log(2\pi)] + 2[\mathrm{df}(\lambda) + 1] \quad (4)$$

The term $n[1 + \log(2\pi)]$ is constant across models fitted to the same channel and subject, so it does not affect model comparison within a channel. When comparing models across subjects or absolute AIC values are needed, the full expression in Eq. (4) should be retained.

The effective degrees of freedom $\mathrm{df}(\lambda)$ are computed from Eq. (2) using the singular values of the design matrix X and the regularization parameter $\lambda$ selected for that fit. The AIC is computed for each subject, channel, and candidate model (i.e., choice of $\lambda$), enabling information-criterion-based model selection.
